# Structural equation models suggest that on-farm non-crop vegetation removal is not associated with improved food safety outcomes but is linked to impaired water quality

**DOI:** 10.1101/2022.09.19.508625

**Authors:** Daniel L. Weller, Tanzy M.T. Love, Donald E. Weller, Claire M. Murphy, Brian G. Rahm, Martin Wiedmann

## Abstract

While growers have reported pressures to minimize wildlife intrusion into produce fields through non-crop vegetation (NCV) removal, NCV provides key ecosystem services. To model food safety and environmental tradeoffs associated with NCV removal, published and publicly available food safety and water quality data from the Northeastern USA were obtained. Because data on NCV removal are not widely available, forest-wetland cover was used as a proxy, consistent with previous studies. Structural equation models (SEMs) were used to quantify the effect of forest-wetland cover on (i) food safety outcomes (e.g., detecting pathogens in soil) and (ii) water quality (e.g., nutrient levels). Based on the SEMs, NCV was not associated with or had a protective effect on food safety outcomes (more NCV was associated with a reduced likelihood of pathogen detection). The probabilities of detecting *Listeria* spp. in soil (Effect Estimate [EE]=-0.17; *P*=0.005) and enterohemorrhagic *Escherichia coli* in stream samples (EE=-0.27; *P<*0.001) were negatively associated with the amount of NCV surrounding the sampling site. Higher amounts of NCV were also associated with lower nutrient, salinity, and sediment levels and higher dissolved oxygen levels. Total phosphorous levels were negatively associated with the amount of NCV in the upstream watershed (EE=-0.27; *P*<0.001). Similar negative associations (*P<0*.*05*) were observed for other physicochemical parameters, such as nitrate (EE=-0.38). Our findings suggest that NCV should not be considered an inherent produce safety risk or result in farm audit demerits. This study also provides a framework for evaluating environmental trade-offs associated with using specific preharvest food safety strategies.

**Importance:** Currently, on-farm food safety decisions are typically made independently of conservation considerations, often with detrimental impacts on agroecosystems. Co-managing agricultural environments to simultaneously meet conservation and food safety aims is complicated because farms are closely linked to surrounding environments, and management decisions can have unexpected environmental, economic, and food safety consequences. Thus, there is a need for research on the conservation and food safety trade-offs associated with implementing specific preharvest food safety practices. Understanding these trade-offs is critical for developing adaptive co-management strategies and ensuring the short and long-term safety, sustainability, and profitability of agricultural systems. This study quantifies tradeoffs and synergies between food safety and environmental aims, and outlines a framework for modeling trade-offs and synergies between management aims that can be used to support future co-management research.

## Introduction

Currently, produce growers face pressure from external, non-regulatory groups (e.g., buyers, auditors) to meet a growing number of food safety aims, with potential economic and environmental costs (1–6). Many of the specific food safety practices being recommended by these external, non-regulatory groups were developed without considering conservation or other farm management objectives, and may run contrary to them (1, 3, 7–9). For example, a 2006 *Escherichia coli* O157:H7 outbreak linked to spinach was putatively attributed to wild boar intrusion into Salinas Valley, CA, USA produce fields, although other sources (e.g., irrigation water) were also proposed (10). Following this outbreak, auditors and buyers pressured growers to adopt practices they assumed would reduce intrusion by wildlife that may carry foodborne pathogens into produce fields. One such practice was the removal of on-farm wildlife habitat, including non-crop vegetation [e.g., riparian buffers; (3, 7)]. A 2007 survey of California growers reported that auditors told 19% and 39% of growers to remove on-farm non-crop vegetation and wildlife, respectively, while 10% reporting losing points on audits because of on-farm non-crop vegetation; these percentages were even higher among leafy greens growers (1). Gennett et al. (7) compared land cover in the Salinas Valley before and after the 2006 outbreak, and found that 13% of riparian and wetland vegetation was converted to bare ground or crops, or otherwise degraded, and that 8% of wildlife corridor vegetation was lost. They also evaluated a proposal from corporate buyers for 120 m bare ground buffers around fields, and found that establishing a 120-m buffer would remove 203,132 hectares of non-crop vegetation across 45 California counties, including >20% of riparian and wetland habitat in 12 counties (7). Despite the potentially adverse environmental effects of removing non-crop vegetation, there is limited and conflicting evidence to suggest that removing non-crop vegetation improves on-farm food safety or reduces wildlife intrusion. While several studies have associated proximity to non-crop vegetation with an increased likelihood of detecting pathogens in field soils [e.g., (11, 12)], distance to non-crop vegetation was strongly correlated with other potential risk factors, such as distance to roads and livestock operations in these studies. Conversely, several studies that examined the relationship between foodborne pathogen presence in pre-harvest samples and on-farm non-crop vegetation failed to find evidence of an association [e.g., (13–15)].

Non-crop vegetation provides critical ecosystem services [e.g., erosion prevention, water filtration; (16– 21)], so its removal degrades environmental health and the economic resiliency of farm communities. Riparian, vegetative, and forested buffers mitigate the environmental impacts of farm operations as buffers can reduce the amount of pollutants (e.g., nitrogen, phosphorus, sediment, manure) in farm runoff that reaches streams, and can improve stream habitat by providing shade (18, 19, 22–24). For example, a meta-analysis that compared nitrate removal efficiencies for different types of vegetated buffers found that mean removal efficiencies ranged between 54% (herbaceous) and 85% (forested wetland) (19). Multiple studies have also shown that fecal contamination of surface water by run-off can be reduced by placing vegetative buffers between fecal sources and waterways (22–24). Several studies further suggested that removing non-crop vegetation may degrade on-farm food safety by favoring wildlife vectors of foodborne pathogens and disfavoring coprophagic organisms that reduce foodborne pathogen prevalence (3, 13, 25–27).

Currently, food safety and conservation can represent conflicting aims for growers (7, 28–30). However, there is limited information on tradeoffs and synergies between conservation and on-farm food safety aims that growers can use to guide management decisions. Quantifying these tradeoffs is complicated by the diversity and mutability of agricultural environments, the complexity and interconnectedness of the processes that impact on-farm food safety and environmental outcomes, and a lack of data for modeling. Since almost all research on the food safety and environmental consequences of on-farm, non-crop vegetation removal has been conducted in California, USA, and the Pacific Northwest (7, 14, 15, 21, 25), there is also a need for research conducted in other produce-growing regions. The present study outlines a conceptual framework that can be used to address this need and to quantify tradeoffs and synergies between on-farm food safety and environmental aims. Specifically, this study focuses on a specific, widely-used food safety practice, the removal or maintenance of on-farm, non-crop vegetation (1, 3, 4, 31). As the environmental endpoint, this analysis considers the effects of non-crop vegetation removal on physicochemical and microbial surface water quality. As the food safety endpoint, this analysis considers the effects of non-crop vegetation removal on the likelihood of detecting foodborne pathogens in fecal, soil, water, and vegetation samples. “Detection” is used here instead of “presence” to be technically accurate since laboratory methods may fail to detect a pathogen even when that pathogen is present. For example, high levels of competing microorganisms may reduce the likelihood of pathogen detection even if the target pathogen is present. Structural equation modeling (SEM) was used to relate non-crop vegetation removal or maintenance to these endpoints because SEM can quantify the strength and direction of linkages within complex, interconnected systems like farms. The models were implemented for the Northeastern US, a produce-growing region with rich data to support model development. Data were available on water quality outcomes from published studies (32–34) and citizen science datasets, including outcomes with food safety implications [e.g., fecal indicator bacteria (FIB) levels]. Several of the water quality datasets [e.g., (35–37)] overlap spatially and temporally with surveys that tested farm and other environmental samples for foodborne pathogens [e.g., (12, 32, 38–40)]. Since we compiled datasets from multiple sources, it was important to determine if methodological differences between datasets would swamp signals of interest. Thus, while the primary aim of this project was to develop a framework for modeling trade-offs and synergies between food safety and environmental aims and to implement this framework as a proof-of-concept for a single food safety practice, a secondary aim was to characterize how methodological differences between datasets were related to differences in outcomes and to implement approaches that accounted for these differences when implementing the SEMs.

## MATERIALS AND METHODS

### Survey and Stream Datasets

This study used historical data from 16 peer-reviewed papers and two citizen science databases where the original source was willing to share the complete dataset (12, 32–46). Specifically, we compiled data from the Northeastern USA that reported (i) the presence or absence of foodborne pathogens in soil, water, vegetation, or wildlife feces, or (ii) dissolved oxygen, fecal indicator bacteria, chloride, conductivity, nutrient, salinity, or sediment levels in streams. These data represent samples from produce farms, waterways in produce-growing areas, and heterogenous landscapes dominated by non-agricultural land uses. The latter was included to ensure variation in land use in the final datasets and that findings were generalizable to farms in areas not dominated by agricultural land use. It is important to note that the 18 datasets compiled here represent a convenience sample and were identified based on information available to the study authors and the Cornell Water Resources Institute.

Data were divided into two overlapping datasets: (i) data from surveys that collected feces, soil, water, and/or vegetation samples from farms, cities, or parks [“Survey Data”; 6,131 samples; (12, 38–44)], and (ii) data for grab samples collected from streams and rivers [“Stream Data”; 18,388 samples; Fig 1; c]. All Survey Data were generated by the Cornell University Food Safety Lab (FSL). In each study, soil, water, vegetation, and/or wildlife fecal samples were collected from farms, cities, or parks (12, 38–44) and tested for *Listeria* spp., *L. monocytogenes, Salmonella*, and/or pathogenic *E. coli* presence using approximately the same methods for all studies (Table 1). Some of the studies included in the Survey Data also collected water samples from rivers and streams; these were the only data included in the Survey and Stream Datasets.

**Figure 1:**
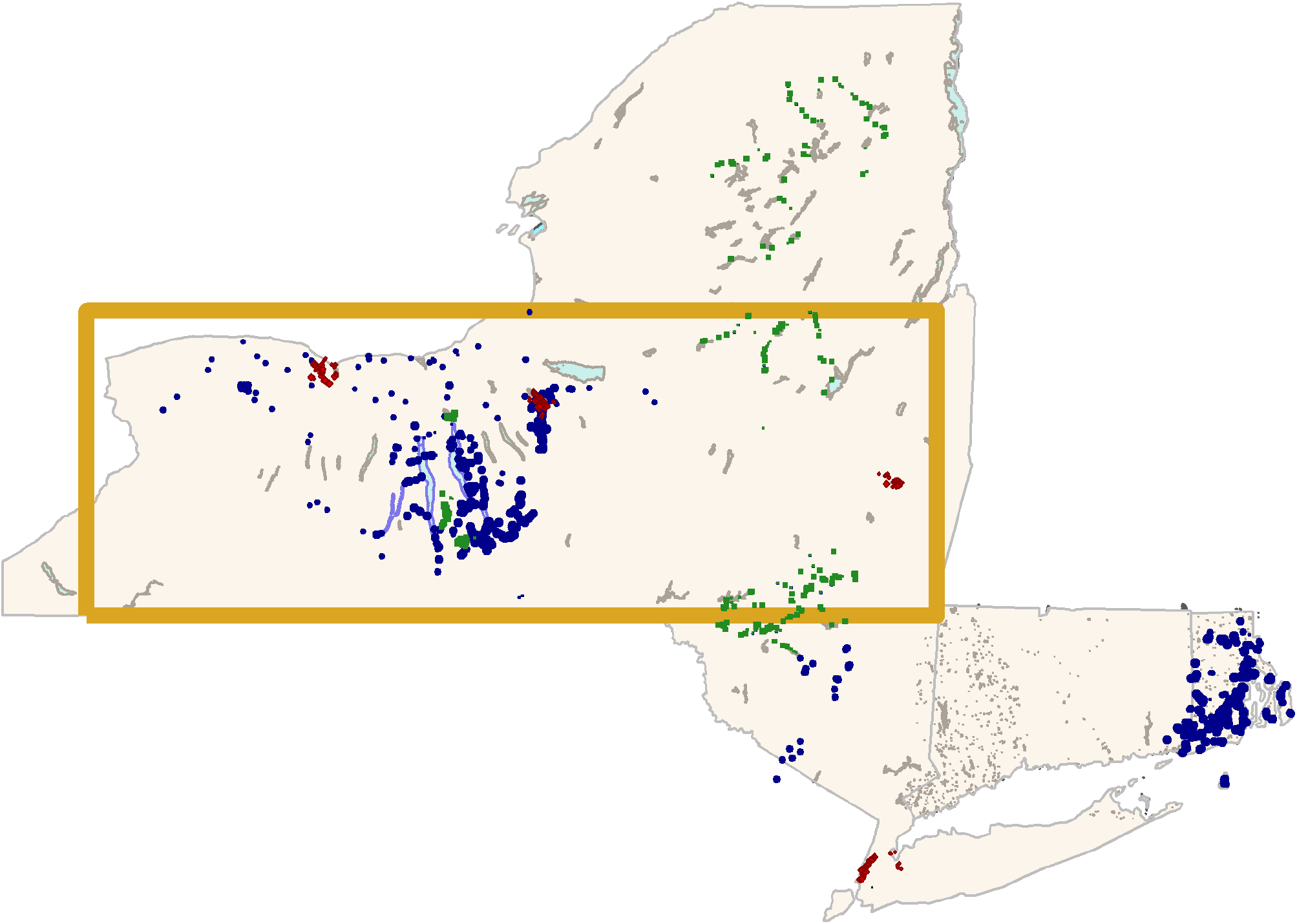
Map of the study region showing all sampling sites in the Stream Dataset (blue points) as well as urban (red diamonds) and parkland (green squares) sampling sites in the Survey Dataset. Due to confidentiality concerns, farm sites cannot be shown, so the yellow box demarcates the area where farm sampling occurred.

**Table 1:** Summary of the studies that constitute the Survey Dataset.

| Project | Study Type <sup>b</sup> | Years | Months | Number of |  |  |  |  |  |  |  | Pathogen Prevalence <sup>h</sup> |  |  |  |
| --- | --- | --- | --- | --- | --- | --- | --- | --- | --- | --- | --- | --- | --- | --- | --- |
|  |  |  |  | Farms <sup>c</sup> | Fields <sup>d</sup> | Samples Collected |  |  |  |  | Totals | (No. of Positive Samples) |  |  |  |
|  |  |  |  |  |  | Feces <sup>e</sup> | Soil <sup>g</sup> |  | Veg. <sup>f</sup> | Water <sup>g</sup> |  | *Listeria* spp. | *L. monocytogenes* | EHEC | *Salmonella* |
|  |  |  |  |  |  |  | Drag Swab | Soil |  |  |  |  |  |  |  |
| A (42) | Long. | 2001-2002 | Apr-Nov | 8 | - | - | - | 426 | 431 | 486 | 1,343 | 24% (316) | 4% (53) | - | - |
| B (43) | Long. | 2009-2010 | Mar-Nov | 5 | 15 | 420 | 84 | 84 | - | 84 | 672 | 28% | 8% (57) | < 1% (2) | 1% (7) |
| C (12) | Long. | 2009-2010 | Feb-Oct | 5 | 20 | 61 | 175 | 178 | - | 174 | 588 | 34% (201) | 15% (88) | 3% (16) | 5% (27) |
| D (44) | Cx | 2010 | Jun-Jul | 10 | 40 | 11 | 40 | 40 | - | 33 | 124 | 40% | 33% (28) | 5% (6) | 8% (10) |
| E (38) | Long. | 2013 | Apr-Jun | 1 | 2 | - | 36 | 486 | 162 | 27 | 711 | - | 10% (68) | - | - |
| F (39, 40) | Long. | 2014 | May-Jun | 1 | 2 | 77 | - | 1,092 | 336 | 132 | 1,637 | 17% (283) | 11% (174) | - | - |
| G (11) | Cx | 2014 | Jul-Aug | 4 | - | - | 1,056 | - | - | - | 1,056 | 20% (208) | 12% (128) | - | - |

<sup>a</sup> Tables S1-3 provide descriptive summaries of these data.
<sup>b</sup> Long. = Longitudinal Study; Cx = Cross-sectional Study
<sup>c</sup> All projects except A and B collected samples from farms. Project A collected samples from 4 urban areas and 4 natural areas (e.g., state parks, national forests), while B collected samples from 5 natural areas.
<sup>d</sup> Fields sampled per farm. Samples in Project A were collected over large, non-continuous urban or natural areas and are not divisible into fields. Conversely, Project B divided each natural area into 3 sub-areas, while Projects C to G sampled farm fields. Specifically, Projects C-F selected a subset of fields for sampling on each farm, while Project G collected samples from all fields on a farm, and thus, the number of fields sampled in Project G was farm-dependent.
<sup>e</sup> All projects that collected fecal samples collected feces in or near the sampled fields opportunistically. Projects C, D, and B collected up to 5 feces per field per sampling visit, while Project F collected any feces observed in or near the sampled fields. Up to 10 g of feces were collected per sample; all samples collected by Projects B and F were 10 g.
<sup>f</sup> Two types of soil samples were collected, representing the soil surface (drag swabs) and sub-surface soil (soil samples). While Projects A and F collected 25-g samples from single points within the sampling unit, all other projects collected composite soil samples consisting of five 5-g samples collected within the sampling unit. Composite soil and drag swabs either represented an entire field (e.g., Projects B-D) or a plot within a larger field (e.g., Projects E and G).
<sup>f</sup> Veg.=Composite vegetation samples consisting of either 25 g (Project A) or 10 g (Projects E-F) of material collected systematically within the sampling unit.
<sup>g</sup> 250-mL grab samples filtered through a 0.45 µm filter.
<sup>h</sup> All projects used the same protocol for enterohemorrhagic *E. coli* (EHEC) and *Salmonella* enrichment. It is important to note that all studies included in the survey data used a culture-based method that included PCR confirmation of the presence of the *eaeA* and *stx* genes; as a result, only EHEC was tested for by the survey studies. In comparison, Projects B-G used the same methods for *Listeria* spp. and *L. monocytogenes* enrichment. Project A used different enrichment and plating media.

The Stream Data were collected by six different labs (47, 48). Methodological differences between the studies represented by the Stream Data are summarized in Tables 2-3. One of the most substantial differences between studies was the method used to detect pathogenic *E. coli*. Ten studies (including those represented in both the Survey and Stream Data) used a culture-based approach for pathogenic *E. coli* detection followed by PCR confirmation of *eaeA* and *stx*. Hence these studies only reported if enterohemorrhagic *E. coli* (EHEC) was detected (defined as isolation of *E. coli* that carried both *eaeA* and *stx*). Three studies used culture-independent methods (i.e., a PCR screen) that separately tested for *stx* and *eaeA*; these studies classified samples as (i) positive for Shiga-toxin-producing *E. coli* (STEC) if a sample tested positive for *stx*; (ii) positive for enteropathogenic *E. coli* (EPEC) if a sample tested positive for *eaeA*; or (iii) positive for EHEC if a sample tested positive for both *eaeA* and *stx*.

**Table 2.** Summary of the studies that constitute the Stream Dataset, including the methods of enumeration and/or detection of microbial targets.

| Lab (Citations) | Years | Months | States | No. of |  | Detection (Method)<sup>a b c d e f</sup> |  |  |  |
| --- | --- | --- | --- | --- | --- | --- | --- | --- | --- |
|  |  |  |  | Watersheds<br>(Sites) | Samples | *E. coli* | *FC* | *TC* | *Enterococcus* |
| Community Science<br>Institute [CSI; (37)] | 2002-2020 | Jan-Dec | NY | 33 (250) | 5,696 | MF<br>(5,411) | - | MF<br>(4,066) | - |
| Food Safety Lab (FSL),<br>Cornell University |  |  |  |  |  |  |  |  |  |
| Survey Studies (12, 38–<br>44) | 2001-2015<sup>g</sup> | Feb-Nov | NY | 58 (228) | 424 | - | - | - | - |
| PAWQ (32, 33, 36) | 2017-2018 | Apr-Oct | NY | 68 (74) | 377 | IDEXX<br>(375) | - | IDEXX<br>(375) | - |
| Green Lab, SUNY<br>Environmental Science,<br>and Forestry (34) | 2015 | Jul-Aug | NY | 2 (10) | 29 | - | - | - | - |
| Onondaga Environmental<br>Institute (35) | 2008-2015<sup>g</sup> | Mar-Nov | NY | 8 (224) | 2,816 | MF (281) | MF<br>(2,658) | MF (96) | MF (288) |
| Richardson Lab, Cornell<br>University (45, 46) | 2017-2018 | Jun-Oct | NJ &<br>NY | 5 (27) | 157 | IDEXX<br>(150) | - | - | IDEXX (153) |
| Watershed Watch,<br>University of Rhode Island<br>[URI; (47)] | 1991-2018 | Jan-Dec | CT &<br>RI | 97 (366) | 8,858 | MF<br>(1,330) | MF<br>(4,166) | - | IDEXX<br>(6,724) |

<sup>a</sup> Tables S1-3 provide descriptive summaries of these data.
<sup>b</sup> *Listeria* spp. testing was performed on 424 and 377 samples collected by the FSL Lab as part of the survey and PAWQ studies, respectively. *Listeria* speciation was performed to determine if *L. monocytogenes* was present for 801 of these 802 samples. While both studies used the same culture-based laboratory methods for *Listeria* isolation, the volume of water collected and the filtration method differed. The survey studies filtered 250 mL through a 0.45 µm filter, while the PAWQ study filtered 348 10 L of water through a modified Moore swab. The remaining 29
samples were separated into 9 L and 1 L aliquots, filtered through a modified Moore swab and a 0.45 µm filter, respectively. These 9 L and 1 L aliquots were treated as separate samples in this study; as a result, there were essentially 830 samples tested for *Listeria* spp., with speciation being performed on 829.
<sup>c</sup> *Salmonella* testing was performed on 191 and 377 samples (N=568) collected by the FSL Lab as part of the survey studies and PAWQ study, respectively, as well as 43 samples collected by the Richardson Lab. While both sets of studies conducted by the FSL used the same laboratory methods to isolate *Salmonella*, the Richardson lab used a culture-independent (i.e., molecular) method. Additionally, the PAWQ *Salmonella* isolation protocol incorporated a PCR screen, which was not part of the FSL survey protocol. For *Salmonella* detection, the PAWQ study collected and filtered 10 L through a modified Moore swab, while (i) the Richardson Lab collected and filtered 10 L using tangential flow ultrafiltration, and (ii) the FSL survey studies collected and filtered 250 mL of water through a 0.45 µm filter.
<sup>d</sup> Pathogenic *E. coli* testing was performed on 191 and 377 samples (N=568) collected by the FSL Lab as part of the survey studies and PAWQ study, respectively, as well as 43 samples collected by the Richardson Lab. The FSL survey studies used a culture-based approach followed by PCR-confirmation using a multiplex PCR, which included testing for the *eaeA* and *stx*. As a result, the FSL survey studies only reported if enterohemorrhagic *E. coli* (EHEC) was detected (i.e., if the isolate had both the *eaeA* and *stx* genes). However, the PAWQ study and Richardson Lab used culture-independent pathogenic *E. coli* detection methods. Both labs used methods that separately tested for the presence of the *stx* and *eaeA* genes, among others. Thus, unlike the FSL survey studies, the Richardson Lab and PAWQ studies were able to identify samples positive for Shiga-toxin-producing *E. coli* (STEC; based on detection of the *stx* gene) and enteropathogenic *E. coli* (EPEC; based on detection of the *eaeA* genes) as well as for EHEC (based on detection of both the *stx* and *eaeA* genes). The sample volumes and filtration methods used for pathogenic *E. coli* detection were consistent with those used for *Salmonella* detection by the respective studies.
<sup>e</sup> One hundred ninety-six and 157 samples collected as part of the PAWQ study and by Richardson Lab, respectively, were tested for the presence of GFD, an avian microbial source tracking marker (MST) marker. Although all studies that performed avian MST detection used the same protocol, the PAWQ used a slightly modified version to improve host specificity (33). One hundred ninety-six, 29, and 157 samples collected as part of the PAWQ study, and by the Green and Richardson Labs, respectively, were tested for HF183, a human MST marker. One hundred ninety-six, 29, and 144 samples collected as part of the PAWQ study, and by the Green and Richardson Labs, respectively, were tested for Rum2Bac, a ruminant MST marker. Forty-three samples collected by the Richardson Lab were also tested for two additional human MST markers (HUMM2 and *B. Theta*), one additional ruminant MST marker (CowM3), and one additional avian MST marker (*Brevibacterium* spp. LA35 16S rRNA-). The water volumes used for MST detection varied between studies. Specifically, 500 mL were used by the Green Lab, 100 mL were used during the PAWQ study, and either 250 mL (N=48 samples) or 100 mL (N=109 samples) were used by the Richardson Lab; all labs filtered samples through 0.45 µm filters.
<sup>f</sup> IDEXX = a proprietary kit used for quantifying levels of specific fecal indicator bacteria as the most probable number of cells present in 100 mL. MF = membrane filtration that quantifies bacterial levels as colony-forming units per 100 mL. For *E. coli*, CSI and URI both used EPA Method 1603 and EPA Method 1604, respectively, while OEI used SM 9222B-2006/SM 9222G-2006 (ELAP 1049). For *Enterococcus*, OEI used SM 9230C-2007 (ELAP 1042), while URI used membrane filtration with mE agar before 2006 but shifted to IDEXX after 2006. For fecal coliforms,
CSI used SM 9222D-2006, and OEI used SM 9222D-2006 (ELAP 1003). URI used a membrane filtration with mTEC agar before 2005 but shifted to IDEXX after 2005. For total coliforms, CSI used EPA Method 1604, and OEI used SM 9222B-2006 (ELAP 1011).
<sup>g</sup> Specifically, samples were collected by the FSL from 2001 to 2002, 2009 to 2011, and 2013 to 2014, and by OEI from 2008 to 2009 and 2012 to 2017. All other labs collected one or more samples annually during the period reported.

### Land Cover Data and Characterization

As data on the maintenance or removal of on-farm non-crop vegetation are not widely available, the proportion of land under non-crop vegetation (i.e., forest-wetland cover) was used as a proxy, which is consistent with previous, peer-reviewed studies (15, 21). Conceptually, this is justified because areas in agricultural landscapes in the Northeastern US with a high proportion of forest-wetland cover are areas where non-crop vegetation has been maintained, while areas with less forest-wetland cover are areas where non-crop vegetation has been removed. The proximity of a sampling site to riparian vegetation was also used as a proxy for maintenance/removal of non-crop vegetation in the SEMs built using the Survey Data. The latter is justified since past studies, such as Gennett et al. (7) and Lowell et al. (2), quantified non-crop vegetation loss in response to food safety pressures and reported substantial decreases in on-farm riparian vegetation.

Forest-wetland cover and other land cover data for 2001, 2004, 2006, 2008, 2011, 2013, and 2016 were downloaded from the USGS National Land Cover Database (NLCD), which offers the highest spatial and temporal resolution dataset for the study area. However, the NLCD’s 30 m^2^ pixels may not capture small patches of non-crop vegetation (e.g., hedgerows, windrows). The NLCD data for the year closest to the sampling date was used for each sample. For the Stream Data, land under agricultural cover (NLCD codes 81-82), forest-wetland cover (codes 41-43 and 90-95), and developed cover (codes 22-24) within 366 m (approx.. 400 yds) of the sampling site was quantified. This distance reflects existing guidance on how far fields and agricultural water sources should be from potential sources of foodborne pathogens [e.g., concentrated animal feeding operations; (49)]. For samples in the Stream Data for which a watershed boundary shapefile was available, the inverse distance weighted (IDW) proportions of land in the upstream watershed and upstream stream corridor were calculated (50). For the Survey Data, the IDW proportion of land around each sampling site under agricultural, developed, and forest-wetland cover was calculated [as described in (51)] for 50, 100, 250, 500, 1,000, 2,000, and 5,000 m buffers. Three land-use variables (IDW proportions of land under agricultural, developed, and forest-wetland cover around the sampling site) were included in SEMs built using the Survey Data SEMs. In comparison, SEMs built using the Stream Data included up to five land-cover variables (proportions of area ≤366 m of each site under agricultural, developed, and forest-wetland cover, and IDW proportions of the upstream watershed and stream corridor under forest-wetland cover). Different approaches to parameterizing land cover variables were used for the Survey and Stream SEMs due to inherent differences between terrestrial and freshwater systems. For example, we calculated land use parameters for are watershed upstream of each sampling site. However, there is no analog to upstream land use for terrestrial systems.

### Random effects (RE) models

RE models were implemented to assess the effect of methodological differences on each outcome of interest, and identify methodological confounders that needed to be included in downstream analyses. Specifically, the variance attributable to methodological differences between studies was compared against variance attributable to spatial and temporal factors.

Up to seven spatial (longitude, latitude, site, waterway, town, county, state, and water type [river or stream]) and five temporal (month, season, number of weeks since Jan. 1^st^, and year) RE were considered (Table S4). Because all the Survey Data were generated using the same or similar methods, only two methodological RE were considered for models implemented using Survey Data; these models either included a RE for (i) dataset, which was defined as the unique identifier for the source dataset, or (ii) sample type (i.e., drag swab, fecal, soil, water or vegetation).

For certain outcomes in the Stream Data, no methodological differences needed to be considered because (i) the same or similar methods were used by all labs to quantify levels of or to detect the given outcome (e.g., fecal coliforms, TSS, turbidity), or (ii) all data for the given outcome were generated by one lab [e.g., salinity, soluble reactive phosphorous (SRP)]. For models implemented using the Stream Data, up to six methodological RE were considered including (i) study, (ii) lab, (iii) sample volume, (iv) filtration method [0.45 um filter, tangential flow, or modified Moore swab], (v) detection method [culture-based or PCR-screen], and/or (vi) enumeration method [membrane filtration or most probable number]; Table S4). Separate regression models were implemented for all possible combinations of dataset, RE, and outcome using the lme4 package (52). The outcomes considered were: (i) the likelihood of detecting enterohemorrhagic *E. coli* (EHEC), enteropathogenic *E. coli* (EPEC), *Listeria* spp., *L. monocytogenes, Salmonella*, Shiga-toxin producing *E. coli* (STEC), and host-specific microbial source tracking (MST) markers for avian, human, and ruminant fecal contamination, (ii) log10 *E. coli, Enterococcus*, fecal coliform, total coliform, chloride, conductivity, nitrate, salinity, SRP, total phosphorous, total suspended solids (TSS), and turbidity levels, and (iii) dissolved oxygen levels. The binomial distribution (logit link) and the bobyqa optimizer were used for models with a binary outcome. The variance attributable to each RE for each outcome was calculated using the MuMin package (53).

### Structural Equation Models (SEM)

SEMs, implemented with the lavaan package (54), were used to quantify the direct and indirect effects of non-crop vegetation removal or maintenance on (i) preharvest food safety and (ii) surface water quality. Before implementing the SEMs, continuous data were centered and scaled, so effect estimates (i.e., beta coefficients) for continuous outcomes should be interpreted as the change (in standard deviations, SD) in the outcome for a one SD change in a continuous covariate. Binary outcomes were converted to continuous variables because lavaan cannot handle discrete outcomes. As a result, the effect estimates from SEMs with binary outcomes should be interpreted as the change in the probability of detecting the microbial target given a one SD change in a continuous covariate.

Because some covariates and outcomes were not measured in some studies or samples, SEMs were implemented using full information maximum likelihood estimation with fixed.x set to FALSE. If SEMs could not be fit as initially conceptualized, they were simplified by (in order): (i) removing collinear, correlated, or constant covariates; (ii) removing covariates with high missingness; and (iii) reducing the number of outcomes in the SEM. For example, because the fecal indicator bacteria (FIB) SEM built using the Stream Data did not converge when *E. coli, Enterococcus*, and total and fecal coliforms were included as outcomes, separate SEMs were fit for *E. coli, Enterococcus*, and total coliforms, and for fecal coliforms. Similarly, for the pathogen SEM built using the Stream Data, the SEM did not converge when EHEC, EPEC, and STEC were included as outcomes; in this case, STEC was dropped as an outcome because there was substantial overlap in STEC-positive and EPEC-positive samples (69% agreement) and no samples were STEC-positive and EPEC-negative.

#### Parameterization of SEMs Implemented Using Survey Data

Separate SEMs were used to quantify the effects of agricultural, developed, and forest-wetland cover on the probability of detecting foodborne pathogens in each sample type [i.e., soil samples (drag swab and subsurface soil samples), vegetation samples (e.g., preharvest produce), water samples, and wildlife fecal samples]. The Survey Data included information on the presence-absence of 3 pathogens (EHEC, *L. monocytogenes*, and *Salmonella*) and one index organism for *L. monocytogenes* (*Listeria* spp.). While all four targets were tested for in all sample types (feces, soil, water, and vegetation), not all samples were tested for all pathogens. As a result, there were four outcomes in the soil SEM (EHEC, *Listeria* spp., *L. monocytogenes*, and *Salmonella* detection), three outcomes in the water and feces SEMs (*Listeria* spp., *L. monocytogenes*, and *Salmonella* detection), and two outcomes in the vegetation SEM (*Listeria* spp. and *L. monocytogenes* detection; Table 1).

For the four SEMs implemented using the Survey Data (see Fig S1 for conceptual models), the likelihood of detecting each target was modeled as a function of IDW proportions of land under agricultural, developed, and forest-wetland covers, the proximity of the sampling site to riparian vegetation, air temperature ≤ 24 h before sample collection, number of weeks since Jan. 1^st^, and year. The number of weeks since Jan. 1^st^ was a proxy for season because lavaan cannot handle categorical variables with more than two levels. In the vegetation SEM, the prevalence of the microbial target in feces, water, and soil samples collected from the same sampling site on the same day as a given vegetation sample (i.e., prevalence in paired samples) was also included as a covariate. Similarly, target prevalence in paired water and soil samples was included in the feces SEM, target prevalence in paired feces and soil samples was included in the water SEM, and target prevalence in paired feces and water samples was included in the soil SEM. Paired prevalence was included because pathogens can move between niches within farm environments [e.g., irrigation can transfer pathogens in water to produce and soil; pathogens in feces and soil can splash onto produce during irrigation or rain events; (39, 55–59)]. Because two soil sample types (surface drag swab and subsurface soil) were collected, sample type was included in the soil SEM, as was available water storage at the sampling site (based on USGS SSURGO data: websoilsurvey.sc.egov.usda.gov). To account for spatiotemporal patterns in land cover (e.g., the North-South trend in forest and agricultural cover in Western New York), paths were also included in all four SEMs to model land cover as a function of latitude, longitude, and year. Covariances among the land-use variables, and between air temperature and the number of weeks since Jan. 1^st^ were also specified. Due to convergence issues, this conceptual model was modified during implementation, and the relationship between temperature and *Salmonella* isolation was removed from the water SEM.

#### Parameterization of SEMs Implemented Using Stream Data

For the stream data, SEMs were used to quantify the direct, indirect, and total effects of land cover on (i) the probability of detecting microbial contaminants, (ii) water quality parameter concentrations, and (iii) the probabilities of dissolved oxygen (DO) levels being above or below 6.5 mg/L (the threshold for healthy aquatic ecosystems) and above or below 4.0 mg/L (the threshold for an ecosystem being considered hypoxic). In total, seven SEMs were fit, including (i) a pathogen SEM, (ii) a host-specific microbial source tracking (MST) markers SEM, (iii) a fecal coliform SEM, (iv) a separate FIB SEM, (v) a DO SEM, (vi) a salinity SEM, and (vii) an SEM for all other physicochemical outcomes. The initial conceptual model for the Stream SEMs is in Fig S2; some models had to be adapted and simplified due to collinearity or convergence issues. Unless otherwise noted, water quality outcomes in each SEM were modeled as a function of (i) the proportions of land ≤366 m from the sampling site under agricultural, developed, and forest-wetland covers, (ii) the IDW proportions of the upstream watershed and stream corridor under forest-wetland cover, (iii) whether or not a rain event (>6 mm) occurred ≤ 24 h before sample collection, (iv) year, (v) weeks since Jan. 1^st^, (vi) water temperature, and (vii) log10 turbidity levels at the time of sample collection. As a result, turbidity was a “cause and effect” variable that occupied a path between the land cover covariates and other water quality outcomes (Fig S2). Thus, the direct, indirect, and total effect of each land cover variable on each outcome, except turbidity, was estimated by accounting for the effect of the land cover variable on turbidity. In the initial conceptual model (Fig S2), each land cover variable was modeled as a function of year, latitude, and longitude to account for spatiotemporal patterns in land use. The initial conceptual model also included (i) covariances between latitude and longitude, (ii) among land cover variables, (iii) rain events and weeks since Jan. 1st, and (iv) temperature and weeks since Jan. 1^st^. If the distributions of continuous variables were skewed, a log10 transformation was used. Other modifications from the initial conceptual model are briefly detailed in the remainder of this paragraph. In addition to log10 turbidity, the water quality outcomes included in the FIB SEM were log10 *E. coli, Enterococcus*, and total coliform levels. The model included covariances between (i) region [New England (i.e., CT and RI) versus NY/PA], and latitude, longitude, and enumeration method, and (ii) between *E. coli* and total coliform levels. Flow condition (base flow versus stormflow) was included as an exogenous variable for turbidity, *E. coli*, and total coliforms. Data on flow conditions were not available for samples with *Enterococcus* data. Region was included as an exogenous variable for turbidity, *E. coli*, and *Enterococcus*, but not total coliforms because total coliform data were not collected in New England. Enumeration method was also included as an exogenous variable for all three FIBs. Since the FIB SEM did not converge when log10 fecal coliform level was included as a water quality outcome, a separate fecal coliform SEM was fit. In this SEM, flow condition but not rain event was included as an exogenous variable because rain data were unavailable for samples with fecal coliform data. As the fecal coliform SEM did not converge when variables for region or forest-wetland cover in the upstream watershed and stream corridor were included, these variables were dropped.

The water quality outcomes in the MST SEM were log10 turbidity and the probabilities of detecting avian, human, and ruminant MST markers. There were no deviations from the initial conceptual model for the MST SEM. The methods used to collect and detect the three MST markers were collinear with year, so the effect of year and methodological differences among studies could not be differentiated. Therefore, year was included as a proxy for differences in laboratory and sampling methods among studies. Other ways of accounting for these differences (e.g., including dummy variables for lab) resulted in an SEM that did not converge or failed to improve model fit.

In the pathogen SEM, the water quality outcomes were log10 turbidity and the probabilities of detecting EHEC, EPEC, *Listeria* spp., *L. monocytogenes*, and *Salmonella*. An exogenous variable was included to indicate if a culture-based method or a PCR screen was used to detect *Salmonella*. The methods used to detect EHEC, EPEC, and *Listeria* were not included because EPEC and *Listeria* detection methods were the same across studies, and EHEC detection method was collinear with year. Similarly, differences in sample volume and sample filtration methods were also collinear with year (Table 2), so year again accounted for these and other unmeasured methodological differences among studies. When the covariance between rain events and weeks since Jan. 1^st^ was included, the model did not converge, so this covariance was dropped from the final pathogen SEM.

The water quality outcomes in the physicochemical water quality (PCQ) SEM were log10 turbidity, log10 chloride, conductivity, nitrate, total and soluble reactive phosphorous (SRP), and total suspended solids (TSS). Flow condition (base versus stormflow) was included as an exogenous variable, but rain event was omitted because the model did not converge when both were included. Region was included as an exogenous variable in pathways where chloride, nitrate, total phosphorous, and TSS levels were outcomes but not in pathways where conductivity and SRP were outcomes. Covariances among region, latitude, and longitude were included in the SEM, as was the covariance of turbidity with TSS and covariances among nitrate, SRP, and total phosphorous. Because the SEM did not converge when dissolved oxygen (DO) or log10 salinity levels were included as outcomes, separate DO and salinity SEMs were fit. In the salinity SEM, the direct effect of the two upstream forest-wetland cover variables, temperature, rain events, and flow conditions on salinity could not be quantified; the indirect effect of the forest-wetland variables and flow conditions on salinity could be quantified through the turbidity-salinity relationship. The water quality outcomes included in the DO SEM were log10 turbidity, DO levels, whether or not the waterway had healthy DO levels (DO levels >6.5 mg/L), and whether or not DO levels were hypoxic (<4.0 mg/L). Flow condition and region were included as exogenous variables for all water quality outcomes in the DO SEM. Covariances among region, latitude, and longitude were included in the DO SEM, as were covariances between region and each land-use variable, and between DO level and whether or not DO levels were healthy (above versus below 6.5 mg/L). The methods used to quantify most physicochemical water quality outcomes were the same or comparable across studies and/or collinear with region and/or year (Table 3). Thus, the effect of methodological differences among studies could not be differentiated from the effects of region and year, so methodological differences were accounted for by including region and/or year in the SEMs. Other ways of accounting for these differences (e.g., including a series of lab dummy variables or individual methods variables) resulted in non-convergence or an un-identified SEM.

**Table 3:** Summary of the methods used to generate microbial and physicochemical water quality data in the Stream Dataset. ^ab^.

| Study | Methods (No. of Samples with Data for Given Target) |  |  |  |  |  |  |  |  |
| --- | --- | --- | --- | --- | --- | --- | --- | --- | --- |
|  | Chloride | Conductivity | Dissolved Oxygen | Nitrate | Phosphorous |  | Salinity | Total Suspended Solids | Turbidity |
|  |  |  |  |  | Soluble Reactive | Total |  |  |  |
| Community Science Institute [CSI; (37)] | SM<sup>a</sup> 4500-CL- C-2011 (4,346) | SM 2510B-2011/SM 2450 D-2011 (5,135) | Lamotte Kit 5860 (620) | SM 4500-NH3/ E-2011 (4,019) | EPA Method 365.3 (4,306) | EPA Method 365.3 (4,393) | - | SM 2450 D-2011 (5,085) | Turbidimeter (4,785) |
| Food Safety Lab (FSL), Cornell University |  |  |  |  |  |  |  |  |  |
| Survey Studies (12, 38–44) | - | -- | - | - | - | - | - | - | - |
| PAWQ (32, 33, 36) | - | Hach HQ40d (377) | Hach HQ40d (377) | - | - | - | - | - | Turbidimeter (377) |
| Green Lab, SUNY Environmental Science, and Forestry (34) | - | YSI 650 MDS (28) | YSI 650 MDS (28) | - | - | - | - | - | YSI 650 MDS (28) |
| Onondaga Environmental Institute (35) | - | YSI 650 MDS (2,596) | YSI 650 MDS (2,540) | Lachat 10-107- | - | SM 4500-P E-99 (375) | YSI 650 MDS (2,175) | SM 2540 D-2011 (650) | YSI 650 MDS (2,079) |
|  |  |  |  | 4-1C<br>(436) |  |  |  |  |  |
| Richardson Lab,<br>Cornell<br>University (45,<br>46) | - | - | - | - | - | - | - | - | Turbidimeter<br>(11) |
| Watershed<br>Watch,<br>University of<br>Rhode Island<br>[URI; (47, 48)] | SM<br>4500-Cl<br>(450) | - | SOP 010<br>(2,731) | SM<br>4500-NO3 F.<br>(1,299) | - | SM 4500-P-F<br>(1,309) | - | SOP 099<br>(140) | - |

<sup>a</sup> Tables S1-3 provide descriptive summaries of these data.
<sup>b</sup> SM = Standard Methods for the Analysis of Water and Wastewater Twentieth Edition 1998, published by American Public Health Association, American Water Works Association, and Water Environment Federation. All SOPs for the Watershed Watch Program are published at https://web.uri.edu/watershedwatch/resources/ and were approved by the US Environmental Protection Agency, New England in June 2005

## RESULTS

### Random Effects (RE) Models

RE models were used to assess the impact of methodological differences between studies and to determine if these differences would swamp signals of interest. If a target concentration or likelihood of detecting a given target was robust to methodological differences, variance attributable to methodological factors should be near 0 and/or lower than the variance attributable to spatial or temporal factors. For the Survey Data, methodological variables (i.e., dataset and sample type) accounted for less than 5% of variance in the likelihood of detecting EHEC, *Listeria* spp., *L. monocytogenes*, and *Salmonella*, indicating that the likelihood of pathogen detection was robust to methodological differences. In comparison, sampling site, a spatial factor, accounted for 50% and 61% of the variance in the likelihood of detecting EHEC and *Salmonella* levels, respectively.

#### Little variance in the likelihood of detecting EPEC, *Listeria* spp., and *L. monocytogenes* was attributable to methodological factors; however, substantial variance in the likelihood of detecting EHEC and *Salmonella* was attributable to methodological variables and/or factors collinear with methodological variables

In the Stream Data, the factor that accounted for the most variance in the likelihood of detecting EHEC was year (71%), which was collinear with three methods variables (dataset, detection method, and sample volume). Dataset and volume each accounted for 66% of variance in the likelihood of detecting EHEC, while detection method accounted for 34% (Table S4). Note that these variances sum up to 237%, indicating that a substantial amount of the variance in EHEC detection was jointly attributable to methodological and non-methodological factors. Disentangling these signals and accounting for each separately in the SEMs would be difficult; as a result, year was used as a proxy to account for both temporal and methodological confounders in the downstream SEMs. For EPEC, dataset and lab each only accounted for 14% of the variance in the likelihood of detection (Table S4); site (42%), waterway (37%), county (36%), and month (32%) instead accounted for the most variance in the likelihood of EPEC detection (Table S4). Methodological factors may account for less variance in EPEC compared to EHEC detection because EHEC detection was performed using either molecular or culture-based methods but EPEC was only detected using PCR screens. The methodological variable that accounted for the most variance in the likelihood of detecting *Listeria* spp. (16%) and *L. monocytogenes* (7%) was sample volume. However, the variance attributable to site was approximately two and six times the variance attributable to sample volume for *Listeria* spp. and *L. monocytogenes*, respectively (Table S4). After site, waterway (26%) and year (22%) accounted for the most variance in the likelihood of detecting *Listeria* spp., while year (33%) and waterway (32%) accounted for the most variance in the likelihood of detecting *L. monocytogenes* (Table S4). The greatest variance in the likelihood of detecting *Salmonella* was attributable to year (33%) and site (29%; Table S4). Of the method variables that differed between datasets for *Salmonella* and were not collinear with year, dataset (23%) and detection method (17%) accounted for the most variance in the likelihood of detecting *Salmonella* (Table S4); detection method was included as an exogenous variable for *Salmonella* in the stream pathogen SEM.

#### More variance in microbial and physicochemical water quality parameter levels was attributable to spatial compared to methodological factors

With regard to non-pathogen-related outcomes in the Stream Data, sampling site was the factor that accounted for the most variance in log10 *E. coli, Enterococcus*, fecal coliform, chloride, conductivity, nitrate, soluble reactive phosphorous (SRP), total phosphorous, salinity, and turbidity levels as well as the likelihood of detecting human and ruminant MST markers. Waterway was the factor that accounted for the most variance in log10 TSS and the likelihood of detecting avian MST markers (Table S4). The variance in water quality parameters attributable to methodological variables was lower than that attributable to spatial variables for all parameters except total coliforms. The variance attributable to site was more than twice that attributable to dataset, laboratory, or other methodological variables for chloride, conductivity, *E. coli*, fecal coliforms, human MST marker detection, total phosphorous, and total suspended solids (Table S4). Because differences in methods for detecting MST markers and enumerating water quality parameters were collinear with region [New England (CT and RI) versus NY and PA] and year, region and year were included as exogenous variables in these SEMs. Other ways of accounting for methodological differences (e.g., including a series of lab dummy variables, including individual methods variables) were considered but generally resulted in SEMs that did not converge or were un-identified. Models that included the method used for FIB enumeration did converge, and due to the large amount of variance in log10 enterococcus (25%) and total coliform levels (15%) attributable to enumeration method (Table S4), enumeration method was included in the FIB SEM.

### Structural Equation Models (SEMs)

#### All SEMs met the criteria for acceptable model fit

Fit statistics for all SEMs were at or near the thresholds for acceptable model fit (i.e., a Comparative Fit Index [CFI] > 0.90; see Table S9). For example, all SEMs had a root mean squared error ≤10, indicating an acceptable fit. Additionally, all SEMs, except the soil SEM built using the Survey data, had a comparative fit index (CFI) ≥0.90, which indicates good fit and means that ≥90% of the covariation in the data could be reproduced with the hypothesized model; the CFI for the soil SEM was 0.86. The fact that some statistics were slightly outside the range indicating good fit (e.g., CFI for the Soil SEM) could be a product of missing data or unmeasured confounding variables. Since this study was conducted to quantify the direct and indirect effect of non-crop removal/maintenance on food safety and water quality outcomes while controlling for known confounders, exploring alternative model specifications was outside the scope of the present study but should be considered for future analyses.

#### In the Survey SEMs, non-crop vegetation removal was not associated with detecting EHEC, *L. monocytogenes*, and *Salmonella* in fecal, soil, water, or vegetation samples

Overall, we failed to find evidence of a positive association between non-crop vegetation and the probability of detecting food safety hazards (EHEC, *L. monocytogenes*, and *Salmonella*) or food safety-relevant indicator organisms (i.e., *Listeria* spp.). Specifically, there were no significant, positive associations (*P>*0.05) between the probability of detecting any microbial target (i.e., *Listeria* spp., *L. monocytogenes, Salmonella*, or EHEC) in fecal (Table S5), soil (Table S6), water (Table S7), or vegetation (Table S8) samples, and the inverse distance weighted proportion of land around a sampling site under forest wetland cover (Fig. 3, S4). The only significant and substantial (i.e., magnitude not near 0) effect of non-crop vegetation was between forest-wetland cover and the probability of detecting *Listeria* spp. (which is an index organism and not a pathogen) in soil (Table S6). For each standard deviation (SD) increase in the IDW proportion of land ≤366 m from the sampling site under forest-wetland cover, the probability of isolating *Listeria* spp. in soil decreased by 0.17 (95% Confidence Interval [CI]=-0.28, -0.05; *P*=0.005).

Distance to riparian vegetation also was not associated (*P*> 0.05) with the probability of detecting EHEC in soil (Table S6), *Listeria* spp. in water (Table S7), *L. monocytogenes* in feces or water (Table S5, S7), or *Salmonella* in feces, soil, or water (Tables S5-S7). While significant negative associations were found between distance to riparian vegetation and the likelihood of detecting *Listeria* spp. in feces and *L. monocytogenes* in soil, the magnitudes of these effects were minimal compared to other covariates (Figure 2; Tables S5-S6). For example, the effect estimates for IDW proportion of land under agricultural cover (effect estimate [EE]=-0.10; 95% CI=-0.19, -0.02) and sample type (drag swab versus soil; EE=-0.07; 95% CI=-0.09, -0.04) on the probability of isolating *L. monocytogenes* from soil were about 5 and 4 times greater, respectively, than the effect of distance to riparian vegetation (EE=-0.02; 95% CI=-0.03, - 0.01; *P*=0.006; Table S6).

**nFigure 2:**
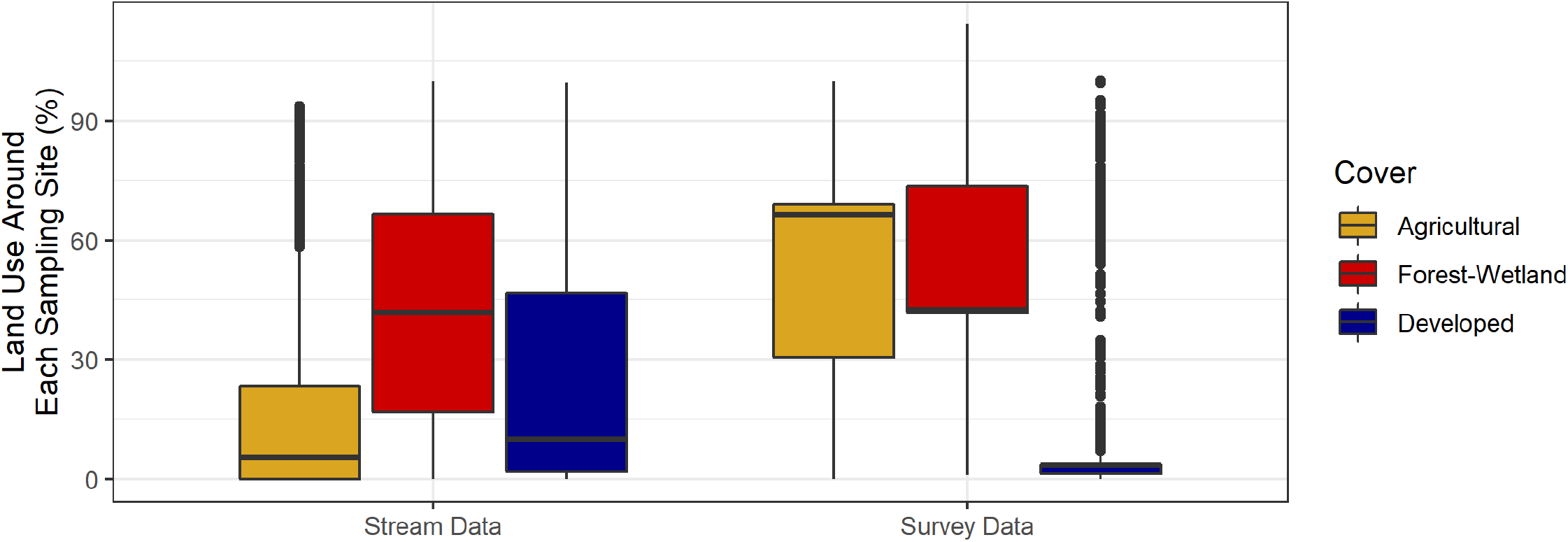
Percent of land around each sampling site in the Stream and Survey Datasets under agricultural, developed, and forest-wetland covers. The radius used for the stream data (1,098 m) was larger than for the survey data (1,000 m) because the stream buffer reflects current guidance on how far animal operations should be from waterways providing irrigation water for leafy greens production (49).

The most consistently significant relationships observed across the four Survey SEMs were between the likelihood of detecting the microbial target and the target’s prevalence in paired samples of other sample types (Figure 3). For example, for each SD increase in the prevalence of *Listeria* spp. in soil samples collected at the same time and location as a fecal sample, the probability of isolating *Listeria* spp. from that fecal sample increased by 0.09 (95% CI=0.06, 0.13; Table S6). Similarly, the probability of isolating *Listeria* spp. and *L. monocytogenes* from vegetation samples increased by 0.10 (95% CI=0.07, 0.14) and 0.04 (95% CI=0.02, 0.05), respectively, for each SD increase in the prevalence of the target in soil collected at the same time as the vegetation sample (Figure 3; Table S8). Similar associations were also observed for the probabilities of isolating *Salmonella* from water (prevalence in soil EE=0.03; 95% CI= 0.00, 0.06) and feces (prevalence in water EE=0.03; 95% CI=0.00, 0.05; Figure 3; Tables S5 and S8). Other factors strongly associated with the likelihood of target detection were air temperature, the number of weeks since Jan. 1^st^, and year (Figure 3;). Temperature (EE=0.05; 95%=0.01, 0.08) and weeks since Jan. 1^st^ (EE=-0.13; 95% CI=-0.22, -0.04) were the only factors significantly associated with the likelihood of detecting EHEC in soil samples (Figure S3).

**Figure 3:**
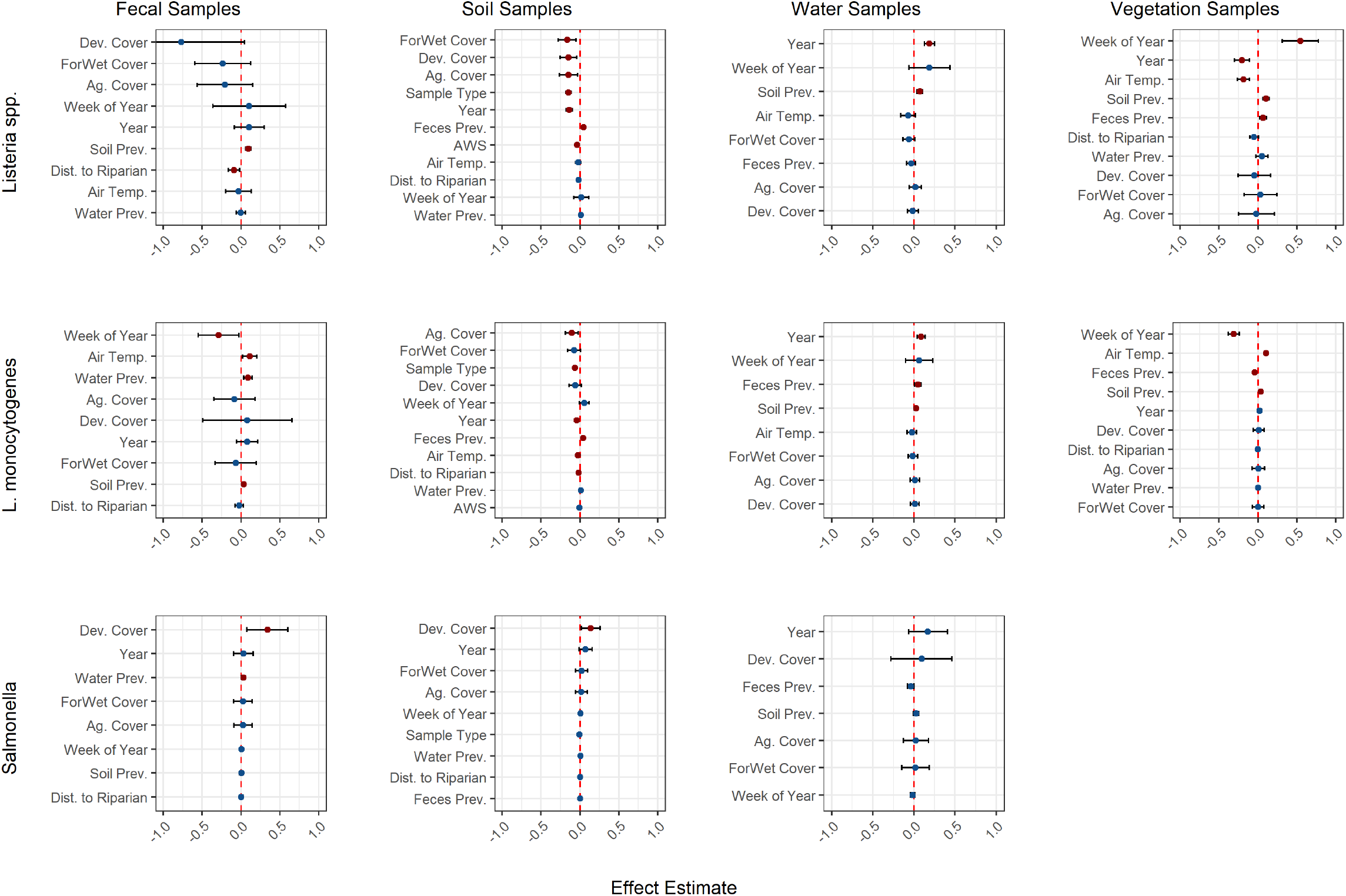
Associations, based on SEMs, between explanatory factors and probability of detecting each microbial target in samples collected as part of the Survey Dataset (red indicates *P*<0.05). Because all continuous factors were centered and scaled, effect estimates can be compared within and across plots, and they should be interpreted as the effect of a one standard deviation increase in a factor on the probability of isolating the target. No vegetation samples were tested for *Salmonella*. See Figure S3 for results where the outcome was the probability of pathogenic *E. coli* in soil samples. The factors along the y-axis were ranked by absolute effect size. Error bars show the 95% confidence interval for the effect estimates. The fit statistics and numerical results for these models are in Tables S5-S9.

#### Non-crop vegetation maintenance was associated with improved microbial and physicochemical water quality

There was no significant association between any forest-wetland cover variable, and fecal coliforms levels or the probability of detecting *Listeria* spp. and *Salmonella* in surface water (*P*> 0.*05* for each of these relationships; see Fig 4-5). Overall, there was a protective effect of non-crop vegetation maintenance on microbial and physicochemical water quality. Specifically, there were significant associations between one or more of the forest-wetland cover variables, and all other microbial outcomes considered (the probability of EHEC, EPEC, *L. monocytogenes*, and avian, human, and ruminant MST marker detection; log10 *E. coli, Enterococcus*, and total coliform levels), as well as multiple physicochemical water quality outcomes (the probabilities of DO levels being >6.5 mg/L or of being <4 mg/L; log10 chloride, conductivity, DO, nitrate, salinity, SRP, total phosphorous, TSS and turbidity levels; Fig 4-5). It, therefore, appears that non-crop vegetation maintenance has a protective effect on microbial and physicochemical water quality--as forest-wetland cover increased, water quality improved.

**Figure 4:**
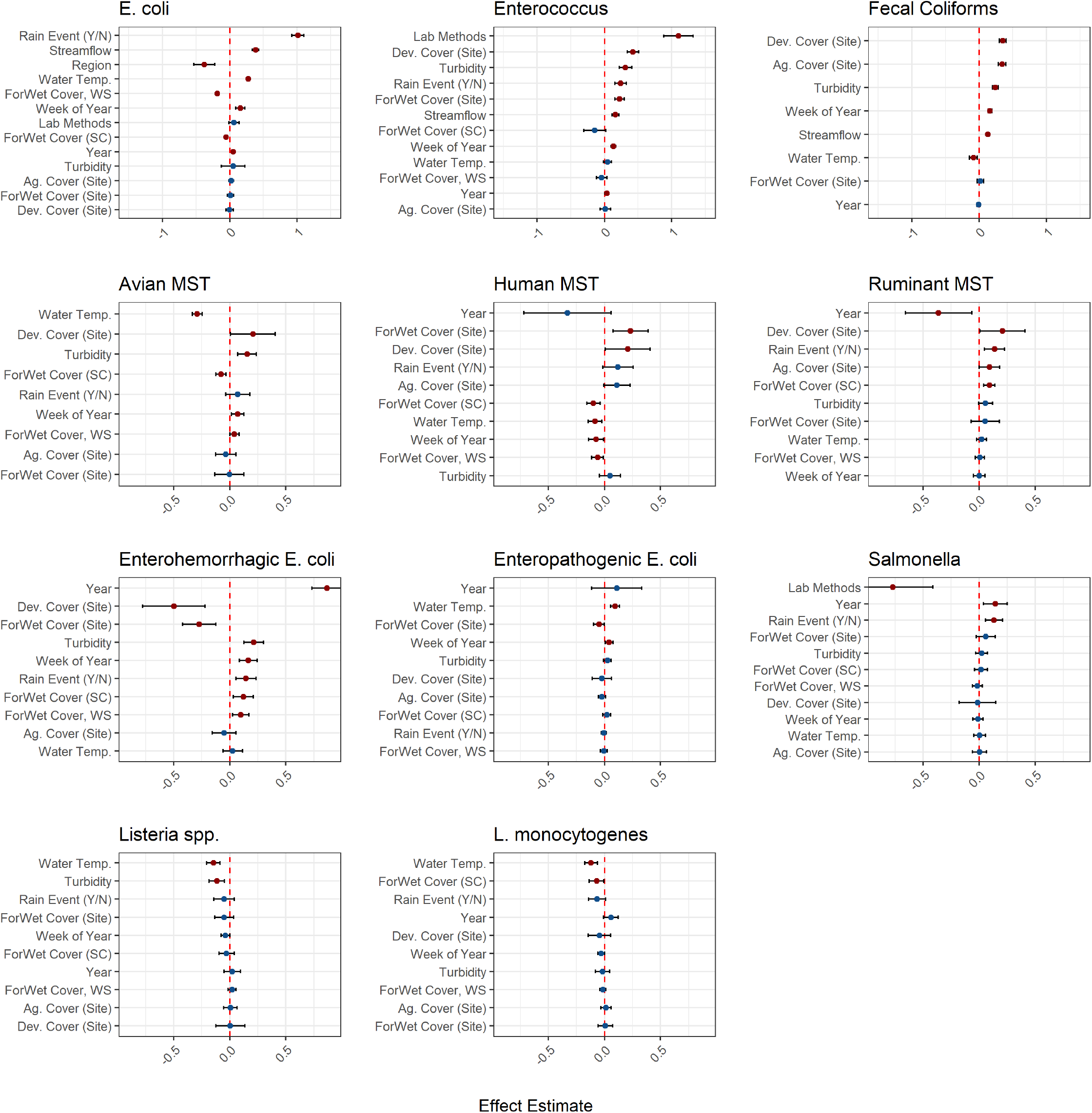
Associations, based on SEMs, between explanatory factors and log10 FIB levels (top-row) or probability of detecting host-specific microbial source tracking (MST) markers (second row), gram-negative pathogens (third row), or gram-positive pathogens and index org-isms for those pathogens (bottom row) in surface water (red indicates *P*<0.05). Because all continuous factors were centered and scaled, effect estimates can be compared within and across plots, and they should be interpreted as the effect of a one standard deviation increase on FIB levels, which were also centered and scaled, or on the probability of detection. The factors along the y-axis were ranked by absolute effect size. Error bars show the 95% confidence interval for the effect estimates. The fit statistics and numerical results for these models are in Tables S9 and S10.

Six water quality outcomes were significantly associated with only one forest-wetland variable. Log10 *Enterococcus* levels and the likelihood of ruminant MST marker detection were positively associated with forest-wetland cover (Fig 4-5), while log10 total coliform and TSS levels, and probability of *L. monocytogenes* and EPEC detection were negatively associated with forest-wetland cover (Fig 4-5).

**Fig 5.**
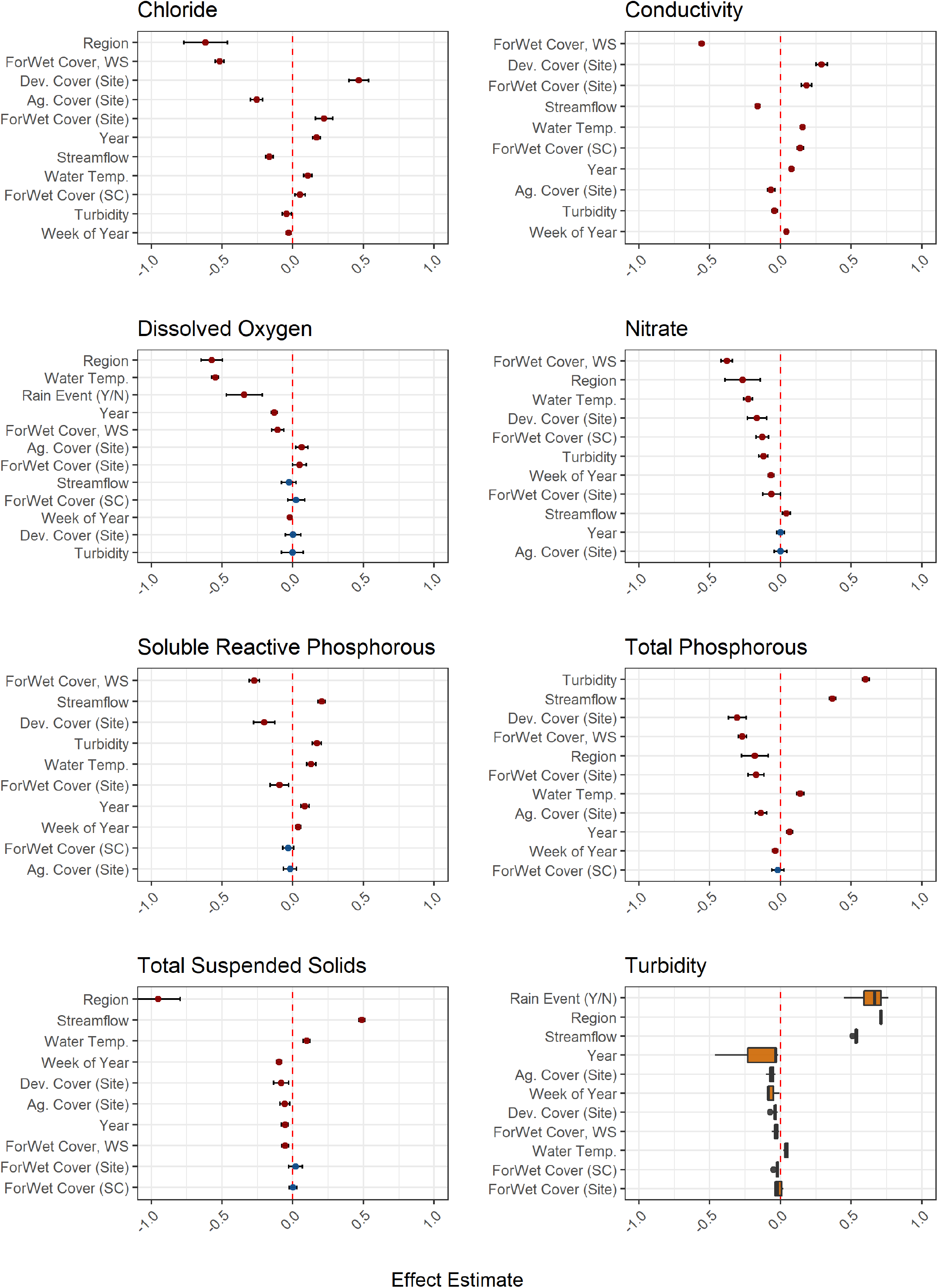
Associations, based on SEMs, between explanatory factors and physicochemical water quality (red indicates *P*<0.05). Associations between explanatory factors with the concentration of total coliforms, salinity, or the probability of DO levels being sufficient to ensure a healthy stream environment (>6.5 mg/L) or hypoxic (<4 mg/L) are shown in Fig. S5. All continuous factors were centered and scaled so effect estimates can be compared within and across plots, and they should be interpreted as the effect of a standard deviation (SD) increase on parameter values, which were also centered and scaled. Before centering and scaling, *E. coli, Enterococcus*, fecal coliform, total coliform, chloride, conductivity, nitrate, salinity, soluble reactive phosphorous, total phosphorous, total suspended solids, and turbidity were log-transformed. Because turbidity was included in all stream SEMs, the distribution of effect estimates for turbidity is shown (as opposed to the point estimates shown in the other figures). Error bars indicate the 95% confidence interval for the effect estimates. The fit statistics and numerical results for these models are in Tables S9 and S10.

Fourteen outcomes were significantly associated with >1 forest-wetland variable. There was a negative association between the water quality outcome and the forest-wetland variable with the largest magnitude of effect for eleven of the 14 (i.e., all nutrient outcomes, all sediment outcomes, and the probability of EHEC detection). There was a positive association between the remaining three water quality outcomes (probability of human MST marker detection, and DO levels being above 6.5 mg/L and above 4.0 mg/L) and the forest-wetland variable with the largest magnitude of effect for the given outcome (Fig 4-5).

Because elevated sediment, nutrient, and microbial levels, and reduced DO levels indicate impaired water quality, these findings suggest that increased forest-wetland cover was associated with improved water quality. This conclusion is also supported by the substantial effect of forest-wetland cover on multiple water quality outcomes, including salinity (EE=-1.28; 95% CI=-1.34, -1.22), conductivity (EE=-0.56; 95% CI=-0.57, -0.54), chloride levels (EE=-0.52; 95% CI=-0.55, -0.49), nitrate levels (EE=-0.38; 95% CI=-0.42, -0.34), SRP (EE=-0.27; 95% CI=-0.31, -0.24), and total phosphorous levels (EE=-0.27; 95% CI=-0.30, -0.24), as well as the probability of EHEC detection (EE=-0.27; 95% CI=-0.42, -0.13), avian MST markers (EE=-0.08; 95% CI=-0.12, -0.03), and EPEC detection (EE=-0.05; 95% CI=-0.10, 0.00), ; Table S10)

#### In addition to non-crop vegetation (forest-wetland) cover, other land cover variables were also associated with water quality outcomes

In addition to the association of non-crop vegetation with water quality, agricultural and developed cover ≤366 m of a sample site were significantly associated with 14 water quality outcomes (Fig 4-5). In Figures 3, 4, and 5, covariates were ranked by the absolute value of their effect; since covariates were centered and scaled before SEM implementation, the relative strength of the association between each covariate and a given outcome can be compared. These figures show that for ten water quality outcomes, the effect of agricultural and/or developed cover was larger than that of forest-wetland cover. The graphs also show that fecal coliform levels and ruminant MST marker detection had the strongest positive association with agricultural cover, while salinity and chloride levels had the strongest negative association. Similarly, chloride, *Enterococcus*, and fecal coliform levels had the strongest positive association with developed cover, while salinity levels and probability of EHEC detection had the strongest negative association (Fig. 4-5). The conflicting results for chloride and salinity are likely an artifact of the fact that salinity was only measured in one study and that this study did not measure chloride (Table 3).

#### The effect of non-crop vegetation variables on water quality outcomes was generally smaller than that of weather variables

For 14 water quality outcomes, the effect of a weather variable (temperature, rain event, or flow condition) was larger than the effect of all forest-wetland cover variables (Fig 4-5). For example, log10 *E. coli* levels decreased by 0.19 (95% CI=-0.21, -0.16) for each 1 SD increase in forest-wetland cover in the upstream watershed but increased by 1.01 SD (95% CI=0.92, 1.10) if sampling occurred ≤24 h after a rain event compared to > 24 h after (Table S10). Log10 *E. coli* levels also increased 0.39 SD (95% CI=0.34, 0.43) if sampling occurred under stormflow as opposed to base flow conditions and 0.27 SD (95% CI=0.25, 0.30) for each SD increase in water temperature (Table S10).

Strong associations between turbidity and multiple water quality outcomes were also observed. For example, total phosphorous levels increased by 0.60 SD (95% CI=0.58, 0.63) for each 1 SD increase in log10 turbidity levels, while salinity increased by 0.22 SD (95% CI=0.18, 0.26), SRP increased by 0.17 SD (95% CI=0.14, 0.02) and the probability of detecting EHEC increased by 0.22 (95% CI =0.13, 0.30) for each 1 SD increase in log10 turbidity levels (Table S10).

#### In addition to non-crop vegetation and weather variables, methodological variables were significantly associated with observed water quality

For 11 water quality outcomes, the effect of methodological variables, including year and region, which were proxies for methodological differences between studies, was larger than the effect of all forest-wetland cover variables (Fig 4-5). Year was significantly associated with 12 of the water quality outcomes and was the covariate with the largest magnitude of effect on the probabilities of detecting ruminant MST markers and EHEC (Fig 4). For EHEC, this is most likely because 250-mL samples were filtered through 0.45 um filters and tested for EHEC using culture-based methods before 2015, but after 2015 10-L samples were filtered using tangential flow or modified Moore swabs and tested for EHEC using PCR-screens. A relationship with region could not be modeled for 13 of the 23 water quality outcomes because these outcomes were measured in NY and PA samples but not in New England samples (Tables 2-3). Of the ten water quality outcomes collected in both regions, all were significantly associated with region (Fig 4-5). Region had the largest magnitude of effect of any covariate on the three DO outcomes, chloride, and TSS (Fig 5).

In addition to year and region, specific methodological variables were explicitly included in the SEMs because they were not collinear with either year or region. Detection method (culture-based versus PCR-screen) was significantly and strongly associated with the probability of detecting *Salmonella*. Similarly, enumeration method was associated with *Enterococcus* and total coliform levels but was not associated with *E. coli* levels (Fig 4-5). Briefly, *Enterococcus* levels were higher when enumerated using membrane filtration compared to IDEXX Enterolert, while total coliform levels were lower when enumerated using membrane filtration compared to IDEXX Colilert (Table S10). *Salmonella* detection was less likely when molecular-based, as opposed to culture-based methods, were used; in interpreting these results, it is important to note that (i) some culture-based methods included a PCR screen, (ii) only one study used molecular methods, and (iii) 13 times as many samples were tested for *Salmonella* using culture-based as opposed to molecular approaches.

## Discussion

This study developed a framework for modeling trade-offs and synergies between food safety and environmental aims. This framework was then implemented as a proof-of-concept to assess how a single food safety practice (removal/maintenance of on-farm non-crop vegetation) affects food safety and environmental outcomes. Importantly, our study provides a methodological blueprint for SEM-based analysis of large datasets collected by different studies and practical pre-harvest food safety insights. Specifically, our findings indicate that, at least in the Northeastern US, non-crop vegetation removal does ot effectively mitigate pre-harvest produce safety risks and may have unintended environmental consequences.

### Non-crop vegetation removal does not effectively mitigate pre-harvest produce safety risks

In our study, on-farm non-crop vegetation was not associated with or had a negligible effect on the likelihood of isolating foodborne pathogens from feces, soil, water, and vegetation (including preharvest produce). In the Survey SEMs, the proportion of on-farm non-crop vegetation only had one significant and substantial association. This was with the likelihood of isolating *Listeria* spp. from soil samples, and this association was negative. Overall, these findings are consistent with the limited research available on the association between non-crop vegetation and foodborne pathogen contamination in farm environments (13, 15, 60, 61). Karp et al. (15) failed to find evidence that EHEC or *Salmonella* were more prevalent in leafy greens samples collected from fields surrounded by non-crop vegetation compared to other cover types in Western North America and Chile. Instead, Karp et al. (15) found that non-crop vegetation removal was associated with increased pathogen prevalence over time. While all samples tested by Karp et al. (15) were collected post-harvest, studies that collected preharvest samples reached similar conclusions (13, 60). For instance, Sellers et al. (13) concluded that the presence of non-crop vegetation was not associated with the prevalence of *Cryptosporidium, Giardia, Salmonella*, or STEC in rodent fecal samples collected from Central California farms. A similar study found that *Campylobacter* prevalence in avian feces was inversely associated with the amount of natural habitat on Western US farms (60). Several studies that sampled Northeastern produce farms (11, 39, 44) did report that the likelihood of *Listeria* spp. and/or *L. monocytogenes* isolation was higher for samples collected closer to non-crop vegetation, such as forest, wetland, and riparian vegetation, compared to samples collected further away. However, proximity to non-crop vegetation was strongly correlated with other environmental factors, including soil characteristics and proximity to other cover types [e.g., roads, pasture; (11, 39, 44)]. The observed associations between proximity to non-crop vegetation, and likelihood of *Listeria* detection may therefore reflect associations between *Listeria* detection and one of these correlated factors, not an association with non-crop vegetation. Thus, although growers have repeatedly reported an increased pressure to remove on-farm non-crop vegetation to manage pre-harvest hazards (1, 3, 4, 7, 31), this guidance is contradicted by the findings of our and other studies (13, 15, 60, 61). Indeed, both the present study and Karp et al. (15) found evidence that maintaining on-farm, non-crop vegetation may improve pre-harvest produce safety outcomes.

Removal of non-crop vegetation to reduce pathogen introduction by wildlife may not have been associated with reduced detection of foodborne pathogens in the present or previous studies (13, 15, 60, 61) because wildlife may not be the primary on-farm source of pathogens. Indeed, past studies have reported (i) a low prevalence of foodborne pathogens in wildlife feces (27, 60, 62), (ii) that produce contamination by wildlife feces was rare (61), and (iii) that pathogen prevalence was higher in fields adjacent to livestock operations compared to other land uses (11, 12, 15, 60). Moreover, the processes that drive on-farm pathogen dispersal and survival are complex, so even if non-crop vegetation removal reduces pathogen introduction by wildlife, removal may increase pathogen introduction through other processes. For instance, non-crop vegetation can act as a windbreak or buffer to prevent air- and waterborne pathogen dispersal (63–66), which are important routes of pathogen movement (39, 67–73). The reported buffering capacity of non-crop vegetation can explain our finding that higher amounts of forest-wetland cover around sampling sites and in the upstream stream corridor were associated with improved microbial water quality.

Because farms are complex ecosystems, non-crop vegetation removal may not be linked to improvements in food safety outcomes for a variety of other reasons as well. For example, non-crop vegetation removal may not discourage wildlife intrusion, or it may only discourage intrusion by certain wildlife species. Similarly, removal might cause the population of certain species to increase (e.g., generalist, non-native species that do well in simplified habitats), or it might increase pathogen loads in wildlife vectors. Each of these possibilities is consistent with current ecological theory and has been observed (27, 60, 61, 78). For example, Smith et al. found that *Campylobacter* was more frequently detected in feces collected from non-native and feedlot foraging birds compared to native birds and birds in more natural areas (60). Similarly, a Florida study found that natural land cover types were associated with reduced shedding of *Salmonella* by white ibises (78). Moreover, the dilution effect is a well-studied phenomenon wherein higher species diversity has a suppressive effect on pathogen prevalence (27). Because the removal of natural vegetation is frequently linked to reduced wildlife diversity (79, 80), the dilution effect may explain some of the results observed here. While multiple processes may be driving the associations between non-crop vegetation and pathogen contamination in the study reported here, identifying such causal relationships is outside the scope of the present study and the data presented here. Thus, there is a need for additional research on the ecology of foodborne pathogens in farm environments, specifically the interplay between non-crop vegetation, wildlife, and foodborne pathogens.

The use of contaminated surface water for irrigation or frost protection is a source of pathogens in farm environments (39, 74–77), and our study reported here linked non-crop removal to microbial contamination of agricultural water. Together, these findings suggest that non-crop vegetation removal can increase the likelihood of introducing pathogens into farm environments during preharvest surface water use for produce production. This indirect effect of non-crop vegetation removal on preharvest produce safety risks is an unexpected and unintended consequence of non-crop removal, and highlights how our analytical approach and conceptual model can be used to identify and avoid unintended consequences when proposing and implementing new preharvest produce safety management practices.

### Non-crop vegetation removal may have unintended water quality consequences

Our data suggest that non-crop vegetation removal may impair physicochemical water quality. In the present study, higher amounts of non-crop vegetation at the sampling site or in the upstream watershed were associated with lower nutrient (e.g., nitrate, phosphorous), salinity, and sediment levels, as well as higher dissolved oxygen levels. These results are consistent with the scientific literature and further support non-crop vegetation maintenance as a beneficial practice. Multiple studies have noted a protective effect of forest, wetland, and other non-crop vegetation on surface water quality [e.g., by reducing soil erosion, preventing the transport of sediment, nutrients, and road salts in run-off to waterbodies (19, 20, 22, 81, 82)]. Reductions in sediment may also reduce microbial transport from terrestrial to aquatic systems because microbes are frequently transported on soil particles (22, 83). A 2014 review that assessed the efficacy of streamside forest buffers for protecting water quality reported median nitrate removal efficiencies between 55% and 89%, and sediment reductions between 65% and 85% (82). An experimental study reported substantial reductions in TSS (29% to 92%), total phosphorous (38% to 93%), total nitrogen (23% to 92%), and *E. coli* (61% to 94%) even though the buffers were quite narrow [between 1.5 and 6.0 m; (22)]. A second experimental study reported removal efficiencies of 97% for sediment, 85% for nitrate, 91% for total phosphorous for mixed woody-grassy riparian buffers (20). Non-crop vegetation also provides a multitude of other ecosystem services (8, 21, 79, 89–94) as supported by reports that non-crop vegetation promotes lower insect pests levels, higher predator diversity, higher rates of pest consumption, and higher levels of pest parasitoids (21, 93). These studies concluded that removing non-crop vegetation would severely impact biological pest control on farms (21, 93). Non-crop removal also reduces natural pollinators (89, 91, 92), making farms more dependent on managed honeybees and thus vulnerable to the collapse of managed honeybee populations (89). Conversely, full pollination services could be provided by native pollinators for farms near natural land cover without reliance on managed honeybees (89). As a result, maintaining and restoring non-crop buffer strips along streams and around fields are already recommended as best practices in water quality regulations and agricultural guidance documents [e.g., (84–88)]. Our findings support this guidance and suggest that on-farm non-crop vegetation removal as a preharvest food safety practice has a net negative impact on farm and farm-adjacent environments.

## Conclusion

This study relied on published or publicly available data, which means that differences among studies (e.g., in sampling and laboratory methods) may have introduced noise that could swamp signals of interest. The approach used here illustrates how one can quantify the variance in each outcome attributable to methodological differences between studies and use these results to inform SEM implementation [e.g., by including methodological variables and other confounders (e.g., year, region) that explained substantial variance in a given outcome as covariates in the SEMs]. This methodological approach and the datasets reported here provide a blueprint that others can use to leverage multiple datasets collected from different regions and with different methodologies to draw broad insights into the food safety and conservation impacts of horticultural practices. In some cases, this approach may provide more rigorous insights than classical meta-analysis approaches, which are regularly used to synthesize data from multiple studies. However, traditional meta-analysis can be leveraged to improve the SEM approach described here by supporting the unbiased and scientific identification of input data for use in the SEM analysis.

The effect of non-crop vegetation on water quality is complex, but our findings suggest that higher amounts of non-crop vegetation were associated with improved microbial and physicochemical water quality. Regarding practical pre-harvest food safety insights, our study indicates that non-crop vegetation removal in the Northeastern US reduced physicochemical surface water quality while having no impact on or even increasing the likelihood of pathogen contamination of agricultural water and farm environments. Our findings are consistent with previous studies that also found that removing on-farm, non-crop vegetation increased on-farm food safety hazards and impaired agricultural waterways (13, 15, 19, 20, 22, 60, 61, 81, 82). These findings highlight the need for science-based approaches for managing preharvest produce safety risks and the potential for unintended consequences from implementing novel management practices in farm environments without proper testing.

This study suggests that on-farm non-crop vegetation should not be considered an inherent produce safety risk and should not result in demerits during farm audits, at least in the Northeastern US. This study also provides a blueprint for understanding the trade-offs and synergies associated with novel food safety practices. Future studies are needed using data collected with standardized methodologies and from other produce-growing regions. This study is presented as a case study and conceptual model on which those future studies can build.

## Acknowledgments

We were grateful for the technical assistance of Laura Strawn and Todd Walters. We are also extremely thankful for the willingness of Edward Michalenko (Onondaga Environmental Institute), Elizabeth Herron (University of Rhode Island Watershed Watch), Hyatt Green (SUNY College of Environmental Science and Forestry), Nathaniel Launer (Community Science Institute), Noah Mark (Community Science Institute), Ruth Richardson (Cornell University), Stephanie Johnson (Onondaga Environmental Institute), and Stephen Penningroth (Community Science Institute) for their willingness to share data and answer questions about their research.

## Data Availability

All data were previously published or are publicly available (see Tables 2 and 3). For data that are not publicly available, data requests should be directed to the corresponding author of the previously published papers.

## Author Contributions

DLW and MW conceived of the project idea and wrote the grant to fund the research. DLW, DEW, and TL designed the study. DLW oversaw the day-to-day aspects of the project and led the data collection and cleaning efforts with assistance from CM. DLW completed all statistical analyses. All authors contributed to manuscript development.

